# Epigenetic investigation of multifocal small intestinal neuroendocrine tumours reveals accelerated ageing of tumours and epigenetic alteration of metabolic genes

**DOI:** 10.1101/2024.12.02.626017

**Authors:** Amy P Webster, Netta Mäkinen, Nana Mensah, Carla Castignani, Elizabeth Larose Cadieux, Ramesh Shivdasani, Pratik Singh, Heli Vaikkinen, Pawan Dhami, Simone Ecker, Matthew Brown, Bethan Rimmer, Stephen Henderson, Javier Herrero, Matthew Suderman, Paul Yousefi, Stephan Beck, Peter Van Loo, Eric Nakakura, Chrissie Thirlwell

**Affiliations:** University of Exeter Medical School, Exeter, UK; Department of Medical Oncology, Dana-Farber Cancer Institute, Boston, Massachusetts, USA; Cancer Program, Broad Institute of Harvard and MIT, Cambridge, Massachusetts, USA; The Francis Crick Institute, London, UK; Achilles Therapeutics plc, London, UK; NIHR Biomedical Research Centre, Kings College London, London, UK; UCL Cancer Institute, London, UK; MRC Integrative Epidemiology Unit at the University of Bristol, Bristol, UK; Population Health Sciences, Bristol Medical School, University of Bristol, Bristol, UK; NIHR Bristol Biomedical Research Centre, University Hospitals Bristol and Weston NHS Foundation Trust and University of Bristol, Bristol, UK; Department of Genetics, The University of Texas MD Anderson Cancer Center, Houston, Texas, USA; Department of Genomic Medicine, The University of Texas MD Anderson Cancer Center, Houston, Texas, USA; Department of Surgery, Division of Surgical Oncology, University of California, San Francisco, USA; University of Bristol Medical School, Bristol, UK

**Keywords:** DNA methylation, Small intestinal neuroendocrine neoplasms, Multifocality, Epigenetic ageing, Tumour metabolism

## Abstract

**Background:** Small intestinal neuroendocrine tumours (SI-NETs) are the most common malignancy of the small intestine and around 50% of patients present in clinic with multifocal disease. Recent investigations into the genomic architecture of multifocal SI-NETs have found evidence that these synchronous primary tumours evolve independently of each other. They also have extremely low mutational burden and few known driver genes, suggesting that epigenetic dysregulation may be driving tumorigenesis. Very little is known about epigenetic gene regulation, metabolism and ageing in these tumours, and how these traits differ across multiple tumours within individual patients.

**Methods:** In this study, we performed the first investigation of genome-wide DNA methylation in multifocal SI-NETs, assessing multiple primary tumours within each patient (n=79 primary tumours from 14 patients) alongside matched metastatic tumours (n=12) and normal intestinal epithelial tissue (n=9). We assessed multifocal SI-NET differential methylation using a novel method, comparing primary tumours with matched normal epithelial tissue and an enterochromaffin-enriched cell line to enrich for tumour-specific effects. This method reduced the identification of ‘false positive’ methylation differences driven by cell composition differences between tumour and normal epithelial tissue. We also assessed tumour ageing using epigenetic clocks and applied metabolic predictors in the dataset to assess methylation variation across key metabolic genes.

**Results:** We have identified 12,392 tumour-specific differentially methylated positions (Bonferroni corrected p<0.05) which were enriched for neural pathways. The expression levels of the genes associated with top sites were also found to be significantly altered in SI-NETs. Age acceleration was observed across SI-NETs and a variability in epigenetic ‘age’ of tumours within each patient, which we believe is reflecting the ‘order’ in which tumours have developed. This is supported by the correlation of age acceleration with somatic mutational count in the tumours. We have identified SI-NET associated alterations to the methylation patterns in key metabolic genes compared to matched normal tissue, which is more pronounced in metastatic tumours and tumours harbouring chromosome 18 loss of heterozygosity, indicating metabolic differences in these tumour subtypes.

**Conclusions:** We have identified accelerated ageing and changes to regulation of metabolic genes, alongside an epigenetic signature of multifocal SI-NETs. These findings add to our understanding of multifocal SI-NET biology and their molecular differences which may be instrumental in the development of these elusive tumours.

## Background

Neuroendocrine tumours (NETs) develop from cells of the nervous and endocrine systems and can affect multiple tissues across the body. The most common form of NETs affect the small intestine, accounting for up to 18% of NET diagnoses (*1*). Though they are relatively rare, they are the most commonly diagnosed malignancy of the small intestine (*2*), and their incidence is rising globally (*3, 4*) making understanding their molecular biology a research priority. Approximately half of patients diagnosed with small intestinal NETs (SI-NETs) present in clinic with multifocal tumours (*5–7*). These synchronous primary tumours were initially thought to be clonal, however recent genetic studies have indicated they are not (*8–10*). Genomic mapping of multifocal SI-NETs using whole genome sequencing (WGS) has now found that on average just 0.08% of somatic single nucleotide variants (SNVs) and/or indels were shared between primary tumours within a patient (*11*). This study also confirmed a lack of shared driver genes in multifocal SI-NETs, indicating that while genomic alterations play a role in SI-NET pathogenesis, other mechanisms are also needed to drive the development of these tumours.

SI-NETs have an unusually low mutational burden compared with other solid tumours (*12*) and the majority of SI-NETs do not harbour known driver gene mutations (*13*). The most common genetic alterations in SI-NETs are loss of heterozygosity (LOH) of chromosome 18 which occurs in approximately 70% of SI-NETs (*8, 14–17*) and mutations in *CDKN1B* (*9, 18*) which occur in around 8% of SI-NETs (*18, 19*). This low incidence of genetic aberrations driving tumorigenesis makes the epigenetic component of their underlying molecular biology of particular interest in SI-NETs. Disruption of epigenetic modifications such as DNA methylation can lead to altered gene expression, driving tumorigenesis and evolution of these tumours in the absence of driver gene mutations.

Previous investigations into unifocal SI-NETs have identified epigenetic subgroups relating to chromosome 18 LOH status (*20*). More recently, these subgroups were investigated using a multi-omics approach in which tumours with chromosome 18 LOH were found to have lower overall methylation, and 12 genes were identified as having differential methylation and higher expression in this sub-group (*21*). This study also used cell composition estimation from DNA methylation data and identified higher CD14 cell infiltration in tumours with wild type chromosome 18, which corresponded with lower progression-free survival (*21*).

While these methylation differences have improved our understanding of the elusive SI-NET molecular landscape, they have all been identified in cohorts of patients with single SI-NETs, and we continue to have little understanding of the DNA methylation landscape of multifocal SI-NETs. In this study, we investigated the DNA methylation patterns in 100 samples collected from a cohort of 11 patients with multifocal SI-NETs, comparing patterns across multiple primary tumours within a patient, and comparing methylation across molecular subgroups of tumours such as those with and without chromosome 18 LOH. Due to the cell-specificity of DNA methylation patterns, use of an appropriate normal comparison tissue is critical in epigenetic analyses. Previous studies have been hampered by the lack of availability of an appropriate normal tissue comparison for SI-NETs due to the low abundance of enterochromaffin cells in intestinal tissue. To overcome this, we performed initial comparisons with the heterogeneous patient-derived matched normal epithelial tissue and refined this methylation signal using methylation data from an enterochromaffin cell-enriched cell line. This allows triangulation of tumour-specific differentially methylated positions (DMPs), reducing the potential for confounding from cellular heterogeneity in the reference tissue. Further, due to the slow growing nature of SI-NETs, there is currently no method of determining the order in which these multifocal tumours develop. Thus we estimate epigenetic ageing in multifocal SI-NETs using 11 epigenetic clock models to assess if the epigenetic ‘age’ of tumours could be used to determine the order of tumour development.

## Methods

### Patient population

All patients in this study underwent surgery at the University of California San Francisco and were diagnosed with multifocal SI-NETs which had metastasised. In total, 100 samples were collected for this study, including multifocal SI-NETs (n=79), metastases (n=12) and adjacent normal epithelia (n=9) derived from 11 patients (summarised in Figure 1). Each patient included in the study had multiple primary SI-NETs included in the dataset (range 2-16). A full breakdown of samples available for each patient is shown in Supplementary Table 1. Sample collection and DNA extraction for these samples has been described previously (*11*). Each patient provided informed consent in accordance with the protocols approved by the Institutional Review Boards of the Dana-Farber Cancer Institute and the University of California San Francisco.

**Figure 1:**
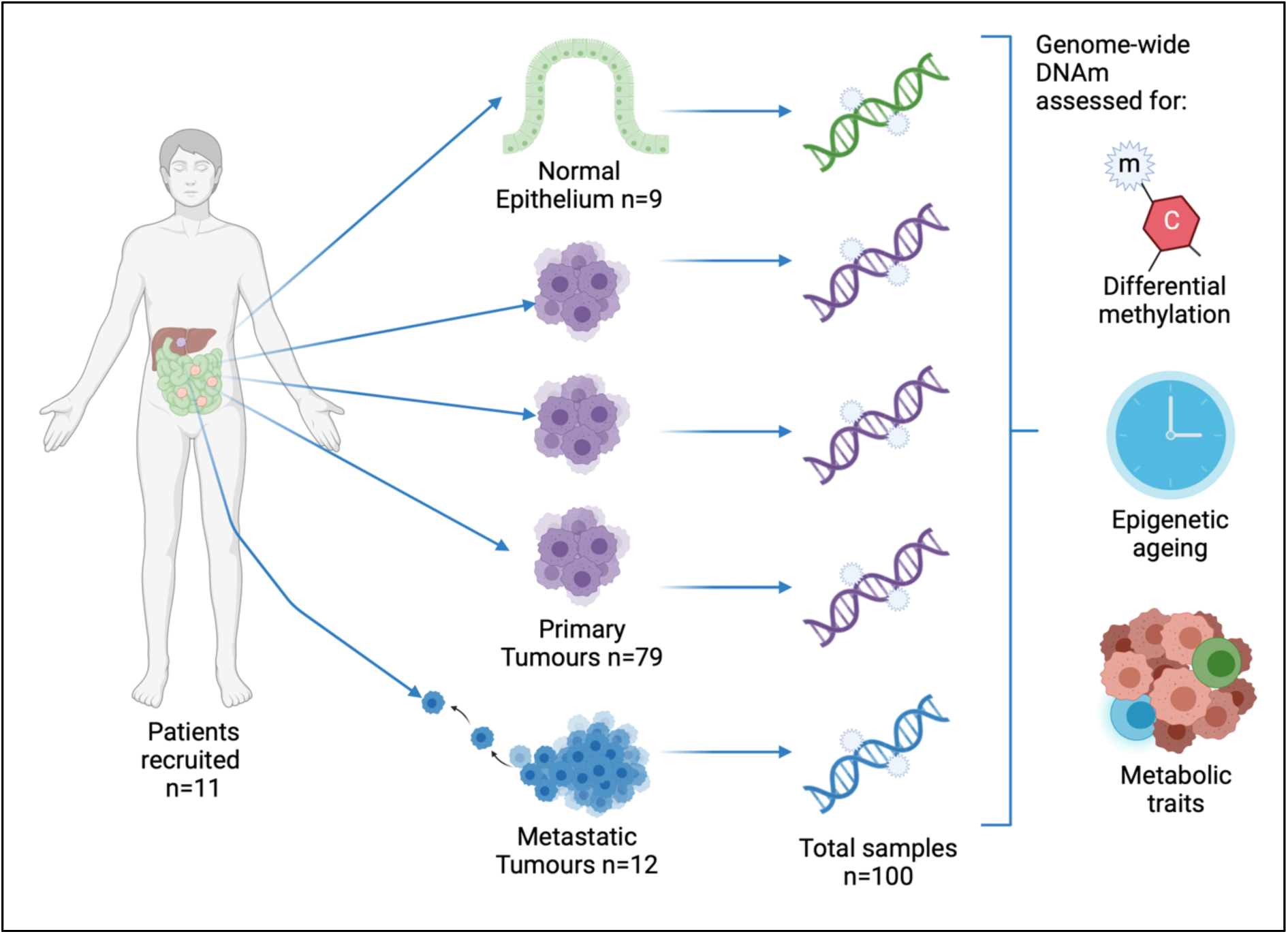
Study design summary. In this study, genome-wide DNA methylation of 100 samples from 11 patients was assessed. Each patient had multiple primary small intestinal neuroendocrine tumours (ranging from 2-16 per patient), and a subset of 9 patients had a matched normal small intestinal epithelial sample assessed. A subset of 8 patients also had metastases originating from their SI-NETs (ranging from 1-2 per patient). DNA methylation data was used to assess differential methylation relating to the multifocal tumours, the epigenetic clock was used to assess the ‘timing’ of tumour development and metabolic traits predicted by DNA methylation were compared between samples.

### Data generation and preparation

Genome-wide DNA methylation microarray data was generated on the Illumina iScan System using the Illumina Infinium MethylationEPIC BeadChip according to the manufacturer’s protocol. For a subset of samples, WGS and RNA-seq data were available on matched tissue. The generation of WGS data has been described previously (*11*). Total RNA was extracted from the samples using RNeasy kit (Qiagen, Germantown, MD). Library construction with ribosomal depletion and sequencing were performed at the Center for Cancer Genomics at Dana-Farber Cancer Institute. Briefly, RNA libraries were sequenced on HiSeq 2500 (Illumina, San Diego, CA) to generate 100-bp paired-end reads. Sequenced reads were aligned to GRCh37 reference assembly using STAR aligner (*22*), and the gene expression levels were estimated with RSEM v1.3.0 (*23*). Samples with less than 5M reads were removed, alongside tumours that clustered with normal intestinal epithelia samples, resulting in matched data for 65 primary tumours and 8 normal samples. Any genes which had a total of less than 10 reads in 8 or more samples were filtered out of the dataset prior to normalisation. Normalisation was performed using DESeq2 (*24*), which uses a median ratios method.

### Overview of data analysis of DNA methylation microarray data

All data analysis was performed in R 4.0.5 (R Development Core Team) using the minfi (*25*), ChAMP (*26*), Enmix (*27*), Limma (*28*) and MissMethyl (*29*) packages. All data was rigorously assessed for quality and presence of batch effects using kernel density plots of methylation beta and M values, alongside principal components analysis and testing of the clustering of the 1000 most variable positions in the dataset using multi-dimensional scaling plots. These measures were taken before and after each step of the analysis to assess the effect of each step during pre-processing. Probe-level and sample-level quality control involved the assessment of detection P threshold of probes. A probe was considered to have failed if it produced a detection P value of >0.01, while a sample was considered to have failed if >5% of probes within that sample had a detection P>0.01 or if they displayed a distorted beta distribution. No samples were considered to have failed these quality measures so data from all samples was retained in the dataset. Both datasets were normalised using BMIQ (*30*) and Noob (*31*) methods to ensure correction of both probe-type and dye-type bias.

### Probe filtering of DNA methylation data

The data was filtered in two ways, to optimise the pre-processing pipelines for downstream analyses:

- For age and metabolic predictions, failed probes (detection P>0.01) were removed along with non-cg probes and probes with a beadcount of less than 3 in >5% of samples. All other probes were retained in the dataset, resulting in 829,090 probes for 100 samples.
- For all other analyses, failed and poor-quality probes were removed as described above, alongside probes mapping to the sex chromosomes. Probes containing known common SNV loci based on the more stringent Zhou 2017 list ‘MASK_snp5_common’ were also removed (*32*), resulting in 661,850 probes for 100 samples.

### Identification of tumour-associated methylation signature

Tumour associated differential methylation was initially assessed through the comparison of the primary tumours with the normal epithelial samples from the cohort. This was performed in the Limma package (*28*) using the ‘block’ method, where each patient was considered one ‘block’ to adjust for relatedness of samples. This resulted in a set of DMPs referred to as the ‘Normal versus Primary Tumour DMPs’ set. All DMPs identified in this study were identified at a Bonferroni P-value significance threshold of 0.05 adjusted for 661,850 tests.

### Use of an enterochromaffin cell-enriched cell line model to refine differential methylation signature

The development of the cell line which is enriched for enterochromaffin (EC) cells has been described previously (*33*). Briefly, biopsies of human intestinal stem cells were cultured as 2d colonies, then committed to the enteroendocrine cell lineage using the EF1a-Neurog3ER-Puro^R^-mCherry construct (*33*). This cell line has been demonstrated to have a high abundance of EC cells and EC cell precursor states at day 5 of incubation, thus DNA was extracted from one sample from this time point, and four bisulphite conversion reactions were run on this sample. Each of these bisulphite-converted samples was run in duplicate or triplicate on the arrays, giving a total of 9 Illumina EPIC array profiles from this time point.

Using the same block method in Limma, DMPs were identified from comparison of the primary tumours and the EC cell-enriched cell line data (‘EC cells versus Primary Tumour DMPs’), and comparison of the cell line data with normal small intestinal epithelial tissue (‘Normal vs EC cells DMPs’). To refine the DMP sites to those with true tumour-related effect, the ‘Normal vs EC cell DMPs’ were subtracted from the ‘EC cells versus Primary Tumour DMPs’. These sites were then tested for overlap with the original ‘Normal versus Primary Tumour DMPs’ set, which resulted in the most stringent set of tumour-specific DMPs.

### Identification of chromosome 18 LOH methylation signature

Differential methylation was assessed between tumours with chromosome 18 LOH and those without. Chromosome 18 LOH status was determined using WGS data from the same samples using a method described previously (*11*). This analysis identified 47 tumours which had full loss of chromosome 18, and three samples which had partial loss or gain of the chromosome. Due to small sample size and lack of power to assess the effect of partial LOH of chromosome 18, the two tumours which had partial loss were combined with the 18 LOH group when assessing methylation, and the sample with partial gain was excluded from this analysis, giving a total of 49 tumours with ‘any’ degree of LOH in chr 18 and 30 tumours with no evidence of chromosome 18 LOH (referred to as chromosome 18 heterozygous tumours). DMP identification was performed using the ‘block’ method in Limma, with each patient being considered one ‘block’.

### Gene Ontology analysis of DMPs

Following the refinement of the significant DMPs using the EC cell-enriched cell line data, the remaining ‘refined’ sites of significance were tested for enrichment of gene pathways using gene ontology analysis. The GOmeth method was used, as implemented in the MissMethyl package (*29, 34*). This method corrects for the bias which can be introduced when performing gene ontology analysis using DNA methylation arrays due to the uneven distribution of probes across the genome (*35*) and the “multi-gene bias” (*34*). The top 20 GO terms were assessed for biological significance and relevance to NET biology.

### Estimation of epigenetic age of SI-NETs

Epigenetic ages of all patient-derived samples were calculated using 11 previously described epigenetic clocks. These included the Horvath multi-tissue clock (*36*), skin and blood clock (*37*), GrimAge clock (*38*), PhenoAge clock (*39*), Hannum clock (*40*), MethylDetectR clock (*41*), epigenetic timer of cancer (EpiTOC) clock (*42*), EpiTOC2 (*43*), HypoClock (*43*), telomere clock (*44*) and the cortical clock (*45*). Epigenetic age acceleration residuals, which is a measure of the residual from a regression of the epigenetic age predictions from each clock on chronological age of the patient, were calculated for the multi-tissue, skin and blood, GrimAge, and PhenoAge estimation methods. These clocks, the telomere clock and the Hannum clock were estimated using the online ‘DNAm Methylation Age Calculator’ (*46*). EpiTOC, EpiTOC2 and the HypoClock were estimated using the R-script ‘*epiTOC2.R’* made available in the EpiTOC2 manuscript (*43*), accessed from https://zenodo.org/records/2632938. The MethylDetectR clock (also referred to as the Zhang clock) was calculated using the ‘MethylDetectR – Calculate Your Scores’ online tool (*47*). The cortical clock was calculated using the freely available R script ‘*CorticalClock.r’*, accessed via the GitHub page https://github.com/gemmashireby/CorticalClock linked in the manuscript (*45*).

The output of all clocks was then assessed in relation to chronological age where appropriate. Matched normal epithelial samples were used to benchmark each clock, assuming these should be closest in age to the chronological age of the patients. Further downstream analyses were performed on the predictions from the skin and blood clock, including testing of the correlation between metastatic tumours and their respective primaries (which was defined using the matched WGS data). The mutational burden of each tumour, as measured from WGS data, was also compared with age acceleration residuals to test for correlation in the two measures.

### Metabolic predictions from SI-NET data

Predictions of metabolic traits were performed on primary tumours, metastatic tumours and matched normal epithelial tissue, using the MethylDetectR (*41*) software. Metabolic traits included body mass index (BMI), body fat, waist-hip ratio (WHR) and HDL cholesterol levels. The MethylDetectR model for metabolic predictions were trained using blood samples from 5,087 people in the ‘Generation Scotland’ study. As this method was developed for application to DNA methylation data derived from blood samples, these models are being assessed as a reflection of metabolic phenotype in the tumours by assessing a composite score of DNA methylation from metabolic genes, rather than for true assessment of BMI, body fat or WHR of patients. In addition to this, CpG sites from the key metabolic genes which are used in these models were assessed for differential methylation between subgroups. These methods were used to compare primary tumours with metastatic tumours and matched normal epithelial samples, and to compare primary tumours with and without chromosome 18 LOH.

## Results

### Dataset overview

The core dataset for this study consisted of 100 samples derived from 11 patients, which were assessed for genome-wide DNA methylation. Multiple primary SI-NET tumours were assessed for each patient (ranging from 2-16 primary tumours per patient, totalling 79 primary tumours). For a subset of 8 patients, metastatic tumours were also assessed (1-2 metastatic tumours per patient, totalling 12 metastatic tumours). For a subset of 9 patients, normal small intestinal epithelial tissue samples were also assessed. Following quality control and pre-processing of the EPIC methylation dataset, 829,090 probes remained in the ageing analyses and 661,850 probes remained for DMP analyses. The beta distributions and Principal Components Analysis clustering indicated the remaining data were good quality and samples primarily clustered by sample type, with normal samples clustering separately from tumour samples as expected (Supplementary Figure 1).

### Differential methylation of multifocal SI-NETs indicates enrichment of neural pathways

The presence of multiple cell types in the normal small intestinal epithelial tissue samples is likely to lead to identification of cell-type related differences which could confound the results and confuse the interpretation of the DMPs identified. To reduce the impact of this confounding we generated DNA methylation array data from a cell line which is enriched for enterochromaffin (EC) cells which we used to refine the methylation analysis and identify ‘tumour-specific’ DMPs. Due to the effect of cell culture and cell composition which are both known to influence DNA methylation signatures, the EC cell-enriched dataset was used for the refinement of DMPs from the initial ‘tumour vs normal’ comparison (summarised in Figure 2).

**Figure 2:**
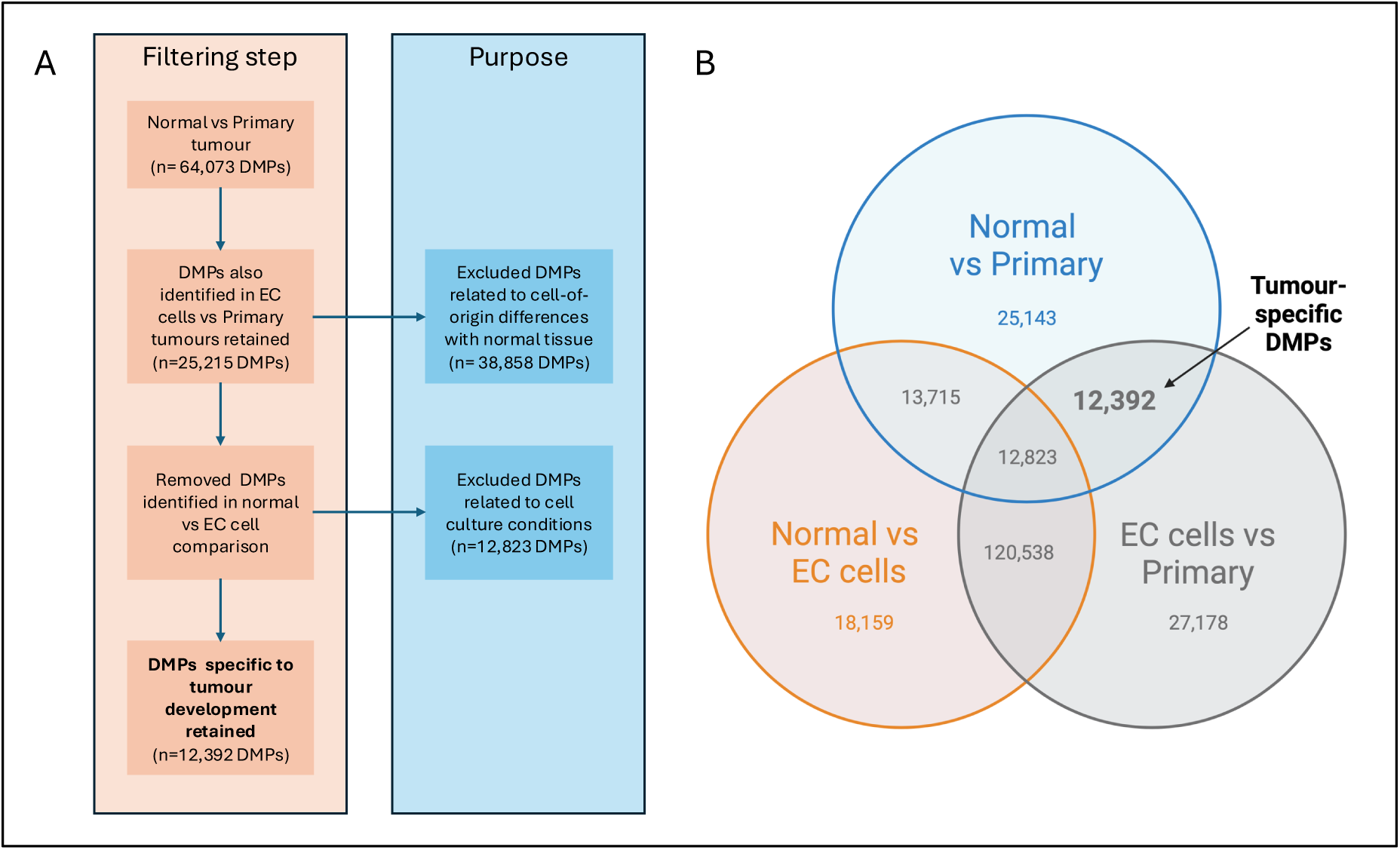
Flow chart and Venn diagram summarising DMPs identified in three comparisons. (A) Flow chart showing filtering steps, summarising the number of probes remaining at each step and the purpose behind each filter in the refinement of tumour-specific DMPs. (B) Three comparisons shown include normal small intestinal epithelia versus primary tumours, EC cell-enriched cell line versus primary tumours and normal small intestinal epithelia versus EC cell line. During analysis the DMPs identified by comparison of primary tumours with both the normal small intestinal epithelia and EC cell-enriched cell line, but not in the comparison of normal small intestinal epithelia versus EC cell-enriched cell line, were considered to be the most biologically relevant to SI-NETs (n=12,392 DMPs). All DMPs indicated in this plot were identified at a Bonferroni-adjusted P<0.05.

Comparison of primary SI-NETs with matched normal epithelial tissue from the small intestine of the same patients identified 64,073 DMPs. While this signature would include tumour associated sites, it is also likely to include false-positive associations driven by differences in cell types present in the patient samples. To reduce the confounding effect of cell type, methylation profiles of primary SI-NETs were also compared with a cell line enriched for EC cells. Comparison of primary tumours with the cell line data (‘EC cells versus Primary Tumour DMPs’) identified a signature of 172,931 DMPs. This methylation signature is likely to include substantial methylation differences introduced by cell culture conditions, and so an additional comparison was made between normal tissue and cell line data (‘Normal vs EC cell DMPs’), which identified 165,235 DMPs. To identify true tumour-related DMPs, the ‘Normal vs EC cell DMPs’ were subtracted from the ‘EC cells versus Primary Tumour DMPs’, leaving 39,570 DMPs which should better reflect the transition from EC cell to SI-NET tumour than comparison with bulk normal intestinal epithelial tissue. This methylation signature is likely to still contain some spurious DMPs caused by cell culture conditions, so an additional step was taken to identify DMPs which overlap with the ‘Normal versus Primary Tumour DMPs’ set. This combination of analyses identified a refined list of 12,392 DMPs (Bonferroni corrected p<0.05) which most accurately reflect the tumour-specific methylation changes in SI-NETs (summarised in Figure 2). Many of the top DMPs mapped to gene bodies and were hypermethylated in tumours compared to matched normal epithelia and EC cell-enriched cell line (Table 1), which reflects activation of these genes in SI-NET tumours, which is supported by the levels of expression of many of these genes in the RNA-seq data (Figure 3). All DMPs which were identified as overlapping between these comparisons were found to be differentially methylated in the same direction (Supplementary Figure 2). Of the refined list of 12,392 SI-NET tumour-associated DMPs identified (Supplementary table 2), 16 mapped to the *SPTBN4* gene (Figure 4) including 4 of the top 10 most significant DMPs identified. The gene expression analyses of matched tissue identified that of the genes associated with the top 20 DMPs, *SPTBN4*, *CACNA1C*, *XKR7*, *ACSL5*, *CEP152*, *DYNC1I1*, *TEAD1* and *CAMTA1* (Figure 3) were significantly differentially expressed (p<0.001).

**Figure 3:**
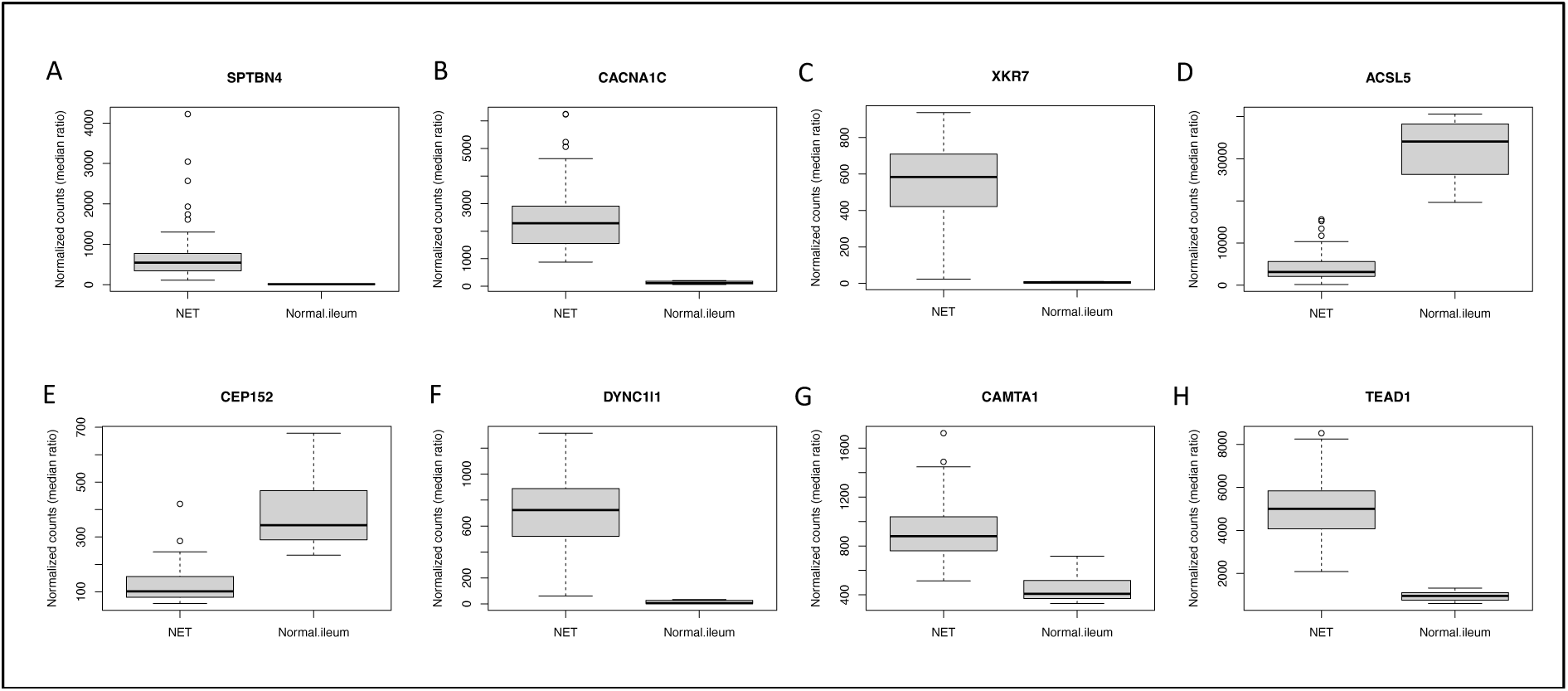
Expression levels of genes associated with top DMPs. The expression levels of 8/10 genes which contained a significant DMP were found to be significantly differentially expressed (p<0.001) between SI-NETs and normal small intestinal epithelial tissue samples.

**Figure 4:**
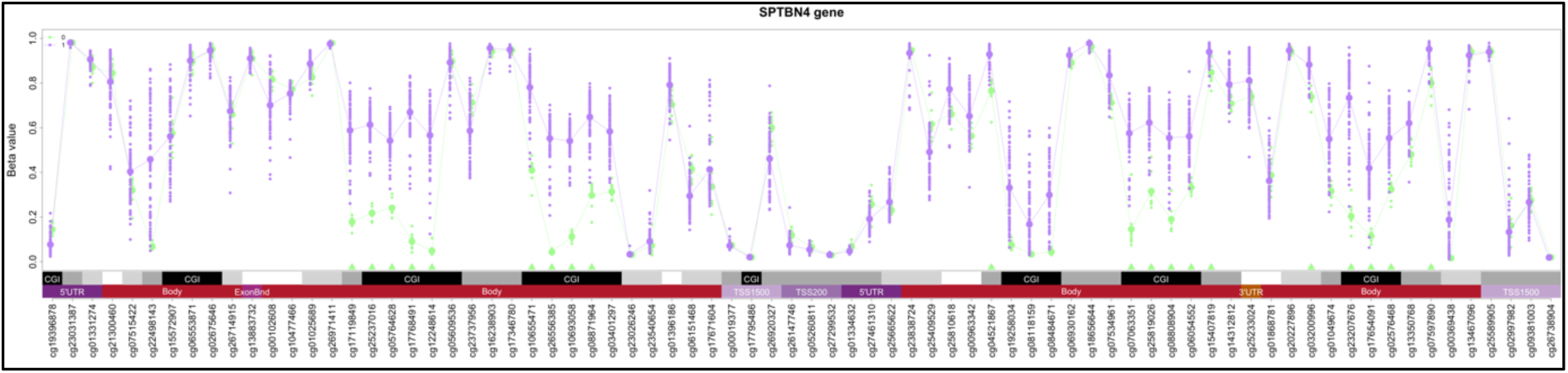
Gene plot of the SPTBN4 gene. Methylation levels of all CpGs included in analysis plotted for primary tumour samples (purple) and matched normal epithelia (green). Tumour-specific DMPs which were identified as significant in the refined analysis (Bonferroni p<0.05) are indicated by green triangles at the base of the plot. Relation of the CpGs to CpG islands is shown below the plot, indicated by black and grey boxes. Relation of the CpGs to genomic elements is demonstrated below the plot, indicating if the CpG maps to the gene body (red), 5’ untranslated region (dark purple), TSS200 (medium purple), TSS1500 (light purple) or 3’UTR (orange).

**Table 1:**
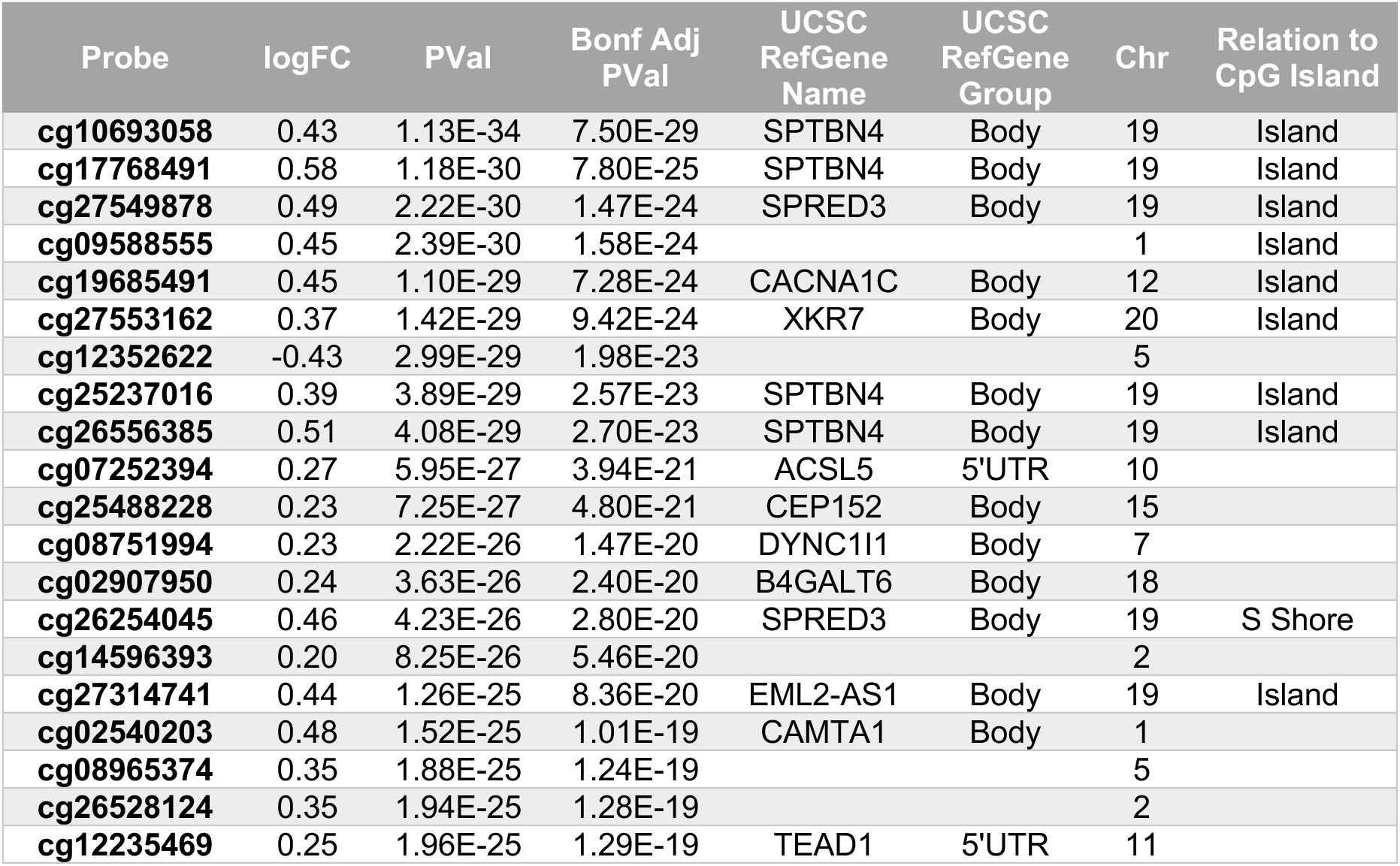
The top 20 tumour-associated DMPs identified in refined analysis. These sites were identified as differentially methylated in SI-NETs when compared with normal small intestinal epithelia and EC cell-enriched cell line, but not in comparison of EC cell-enriched cell line and normal epithelial tissue. The majority of sites (19/20) were hypermethylated in tumours compared to matched normal epithelia and EC cell-enriched cell line. The top DMP maps to the SPTBN4 gene which contains multiple sites of significance across the gene. Chr: chromosome, UCSC RefGene Name/Group: name of gene or gene element annotation as defined by the UCSC Genome Browser, according to the EPIC Illumina manifest.

Pathway analysis of the 12,392 tumour-specific sites identified multiple synapse and neuron-related pathways (summarised in Supplementary Table 3) and several cell signalling and communication related pathways. The top-associated pathway was the ‘synapse’ pathway (FDR p=3.93E-5).

### Epigenetic age predictions suggestive of determining the order of development of multifocal SI-NETs

Epigenetic age was assessed using 11 clocks, to test how the same sample set behaved in a range of different epigenetic age predictors. The multi-tissue clock, PhenoAge clock and telomere clocks are applicable to cancer samples and have been previously tested in other cancer types. EpiTOC, EpiTOC2 and Hypoclock are mitotic clocks which were specifically developed for use in cancer tissue. Hannum’s clock, GrimAge, the skin and blood clock, the MethylDetectR clock and the cortical clock were not developed for use in cancer tissue but were selected due to their wide usage in the field, and/or due to the appropriateness of the tissues for which they were developed. The cortical clock was selected due to the neuronal pathways which are often epigenetically dysregulated in neuroendocrine cells, and the skin and blood clock was used due to its optimisation for epithelial tissue, as intestinal epithelium was used as the ‘control’ sample set in this analysis.

The epigenetic age predictions of all samples varied dramatically between different clocks. In the predictions from the Horvath multi-tissue clock (*36*), the normal samples were consistently predicted to be the youngest, and were systematically younger than the chronological age of the patients (Supplementary Figure 3A). The GrimAge clock (*38*) unexpectedly predicted normal tissue to be consistently the oldest in the dataset (Supplementary Figure 3B) while the PhenoAge clock (*39*) predicted almost all samples to have age deceleration (Supplementary Figure 3C), with no clear pattern shown for normal tissue. The Hannum clock (*40*) predicted both age acceleration and deceleration in tumours, and had no consistent pattern for normal samples (Supplementary Figure 3D). The MethylDetectR clock (*41*) had the lowest variation in age predictions across the samples, and did not display consistent patterns for normal samples. Both EpiTOC (*42*) and EpiTOC2 (*43*) gave extremely similar patterns of predictions (Supplementary Figure 3F and G) however EpiTOC gives a measure of ‘mitotic score’ while EpiTOC2 gives a measure of ‘estimated cumulative stem-cell divisions’. In both of these predictions, the normal samples were estimated to have higher mitotic score/cumulative stem-cell divisions than the tumour samples, which was unexpected and contradicts previous findings (*43*). This higher mitotic score in normal samples was also predicted by the HypoClock (*43*) (Supplementary Figure 3H). One potential explanation for these unexpectedly high predictions is the high division rate of the intestinal epithelium in comparison to tumours. The telomere clock (*44*) predicted telomere length to be shorter in normal samples than in tumours (Supplementary Figure 3I). The cortical clock (*45*) predicted almost all samples to have age acceleration, with the most extreme acceleration seen in normal samples. This clock was applied due to the epigenetic links with neurological pathways in SI-NETs, and as the neuroendocrine cell of origin would be present in very low amounts in the normal tissue used, it is likely less applicable in the normal samples compared to the tumour samples in this study.

In the skin and blood clock predictions (*37*), almost all samples were predicted to have age acceleration (Figure 5A and B) and the normal samples were predicted to be the closest to the actual age of the patients, potentially due to the normal samples being derived from intestinal epithelial tissue. As this clock seemed to perform best in the dataset and had the most interpretable output in terms of defining age acceleration in multifocal tumours, these predictions were used for further analyses. Assessment of the age acceleration differences (Figure 5B) and age acceleration residuals indicated that almost all tumours appear to demonstrate age acceleration. Metastases appeared to have a more extreme age acceleration phenotype than primary tumours, however this was not statistically significant (p>0.05). This trend could be due to ‘older’ tumours having more time to develop metastatic potential or could reflect a more aggressive and accelerated ageing phenotype in tumours which are metastatic. Metastases appeared to cluster with the primary tumour from which they derived from (Supplementary Figure 4), indicating that epigenetic age of the primary tumour is retained when it metastasises, however this effect was not statistically significant (p>0.05), likely due to low power caused by the small size of the dataset. Somatic mutation count including SNVs and insertions/deletions correlated with predicted DNA methylation age (p= 0.0007, Figure 5C). This indicates that SI-NETs which have been growing for longer have more time to develop somatic mutations and emphasises the utility of epigenetic clocks for prediction of the order in which multifocal SI-NET tumours have developed. The variability in epigenetic ages of the tumours also provides further support for the independent evolution theory for multifocal SI-NETs (*8, 11*).

**Figure 5:**
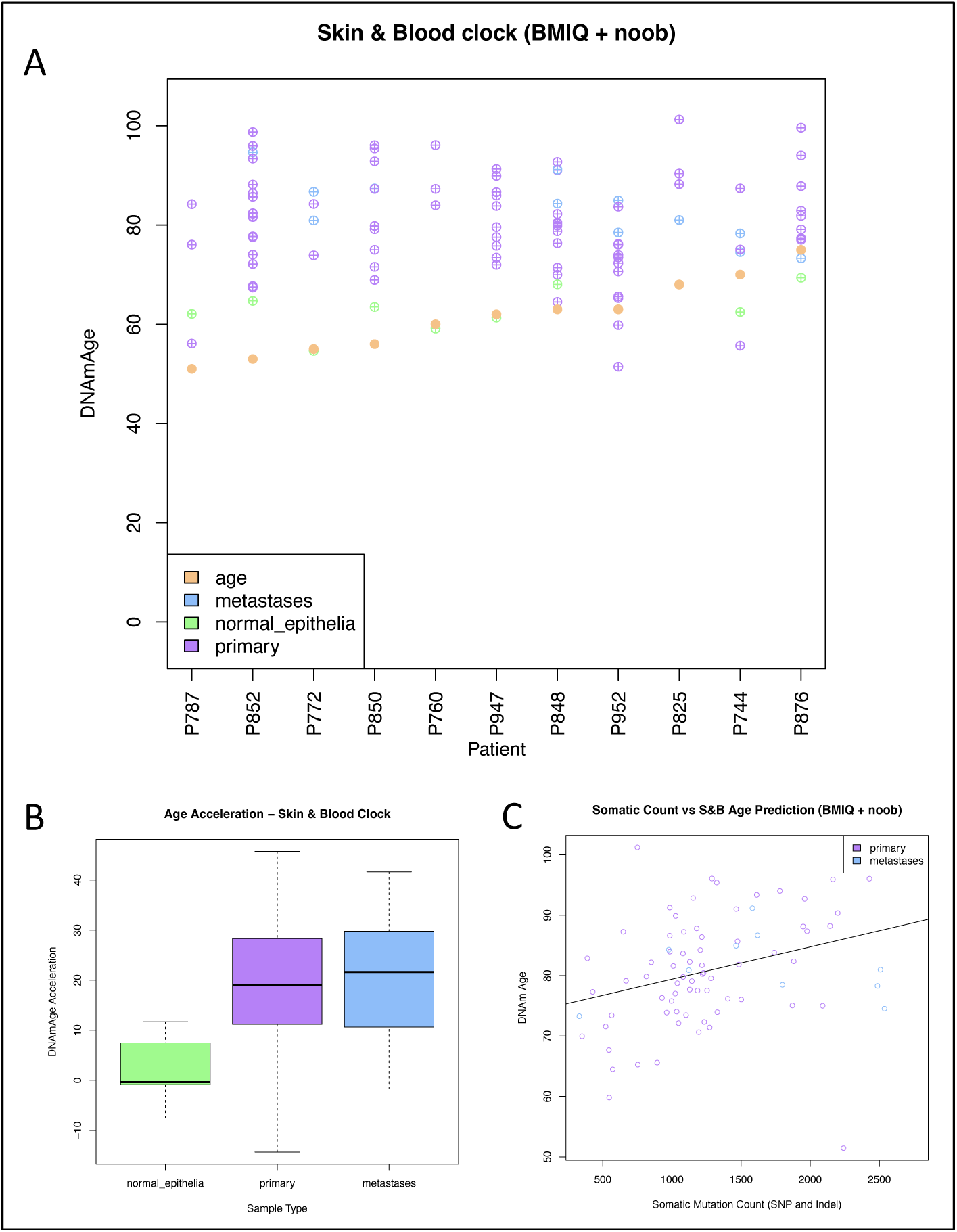
Summary of epigenetic ageing in SI-NET samples using the skin and blood clock. (A) Plot of age predictions for normal epithelia samples (green), primary tumours (purple) and metastatic tumours (blue). Chronological age is indicated with the orange points and patients are indicated in order of chronological age. (B) Boxplot of age acceleration difference for the skin and blood clock predictions. (C) Somatic mutation count from tumours correlates with DNA methylation age in the skin and blood clock (p=0.0007).

### SI-NETs DNA methylation patterns demonstrate altered metabolic phenotype

Very little is known about the metabolic profiles of NETs, though metabolic processes in tumours are known to differ from those in normal tissue. Differences in regulation of gene expression in multiple metabolic pathways can be tested using epigenetic predictors of metabolic traits. To assess the methylation patterns of metabolic pathways, we performed predictions of metabolic traits including body mass index, body fat, waist-hip ratio and HDL cholesterol on SI-NETs, metastatic tumours and matched normal intestinal epithelial tissue. Across all of these predictions, primary tumours demonstrated an altered metabolic phenotype when compared with the normal tissue, and this pattern was more extreme in the metastatic tumours (as demonstrated in Figure 6A). ANOVA tests indicated that the primary tumours, metastatic tumours and normal intestinal epithelial tissue all had significant differences in predicted body mass index (Pr(>F)= 2.45e-12), body fat (Pr(>F)= 7.49e-12, waist-hip ratio (Pr(>F)= 3.46e-05) and HDL cholesterol (Pr(>F)=9.85e-08). The CpG sites which are most informative in driving these predictions demonstrate substantial differences in DNA methylation, as indicated by the differential methylation seen in cg06500161 which is the most influential CpG for HDL cholesterol predictions and second most influential for BMI predictions (Figure 6B). This CpG maps to the *ABCG1* gene which is a key metabolic gene and contained 6 tumour-associated differentially methylated sites in the ‘refined’ DMP analysis described above (Figure 6D). This gene is also found to have differential expression across these groups in matched samples, with higher expression seen in the primary and metastatic tumours compared to normal tissue (Figure 6C). These differences in methylation of key metabolic genes indicate stark differences in metabolism of nutrients within SI-NETs compared to normal tissue which seems to be more drastic in metastatic tumours. This could provide insight into tumour biology, and that metabolic pathways have altered regulation in more aggressive tumours compared to indolent and non-metastatic primary tumours.

**Figure 6:**
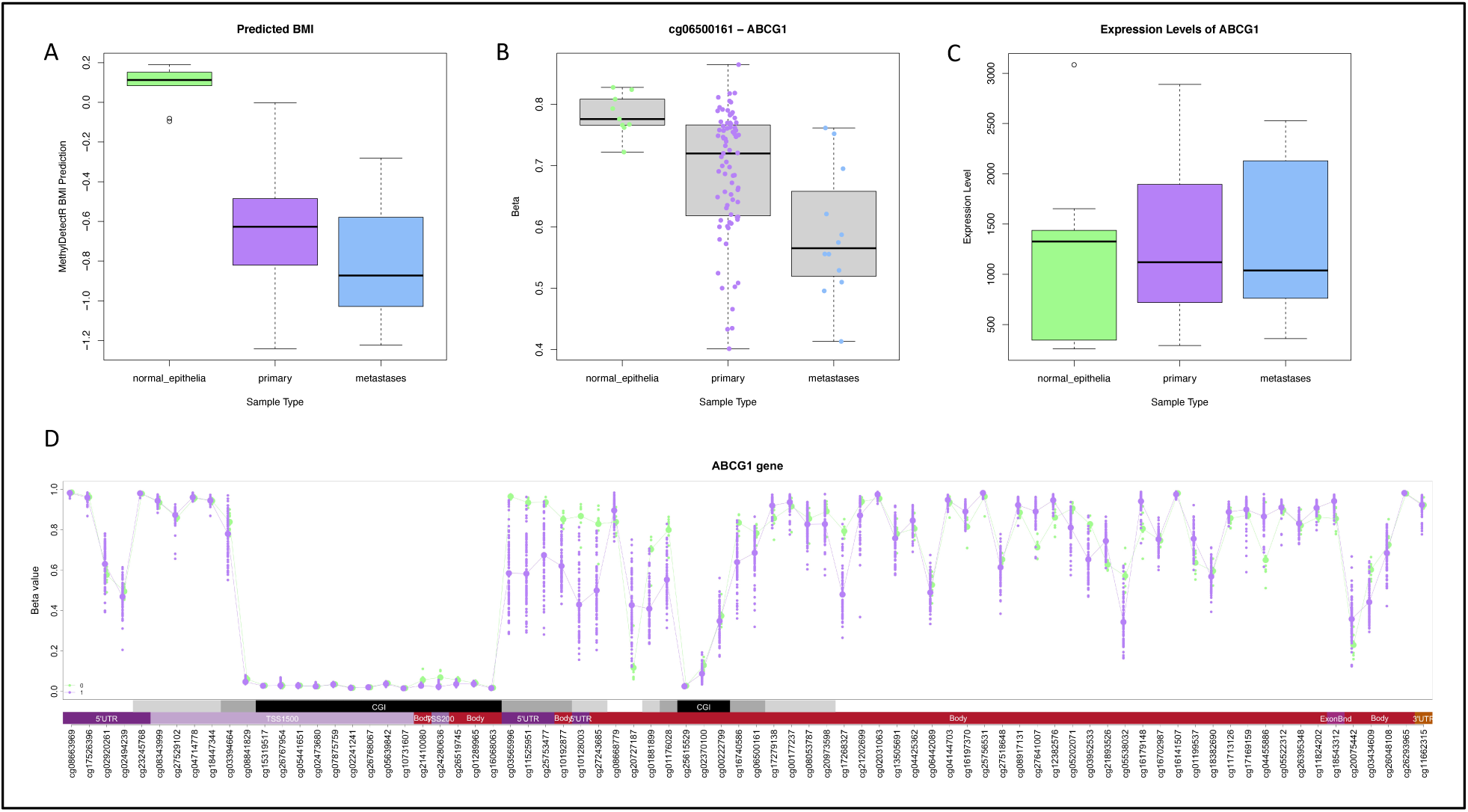
Summary of metabolic trait differences between normal small intestinal epithelia, primary and metastatic tumours. (A) Example of metabolic trait predictions indicating predicted BMI; across all predictions tumours demonstrated an altered metabolic phenotype which was more dramatic in metastatic tumours. (B) Example of differential methylation of a CpG site cg06500161 which is heavily weighted in the metabolic predictors and displays hypomethylation in primary and metastatic tumours compared to normal tissue. This CpG maps to the ABCG1 gene, and gene wide methylation for comparison groups is shown in (D). (C) Gene expression levels of ABCG1 gene; the key metabolic genes which display differential methylation in tumours also appear to have differential expression though it is not as dramatic as the differences seen in methylation levels.

### Tumours with chromosome 18 LOH have distinct methylation signature including disruption of key metabolic genes

To assess the impact of chromosome 18 LOH status on the DNA methylation patterns of this subgroup, we tested for differential methylation signatures between these subgroups and assessed methylation from metabolic pathways using the metabolic predictors. In comparison of SI-NETs with any loss of chromosome 18 (n=49) versus tumours with no loss/wild type chromosome 18 (n=30), 74,593DMPs were identified with FDR adjusted p<0.05, 42 of which reached more stringent Bonferroni significance (Figure 7A). The top 10 DMPs are shown in Table 2. When tested using the metabolic predictors, there was a significant difference in prediction of metabolic markers including predicted BMI and predicted Body Fat (p=0.036 and p=0.00065 respectively), with chromosome 18 LOH tumours having consistently lower predictions (Figure 7B). This reflects differential methylation in the key metabolic genes used in these predictors. This indicates that tumours containing chromosome 18 LOH have a more severe altered metabolic phenotype than tumours with wild type chromosome 18, reflecting potential metabolic differences or differences in nutrient availability in this sub-group.

**Figure 7:**
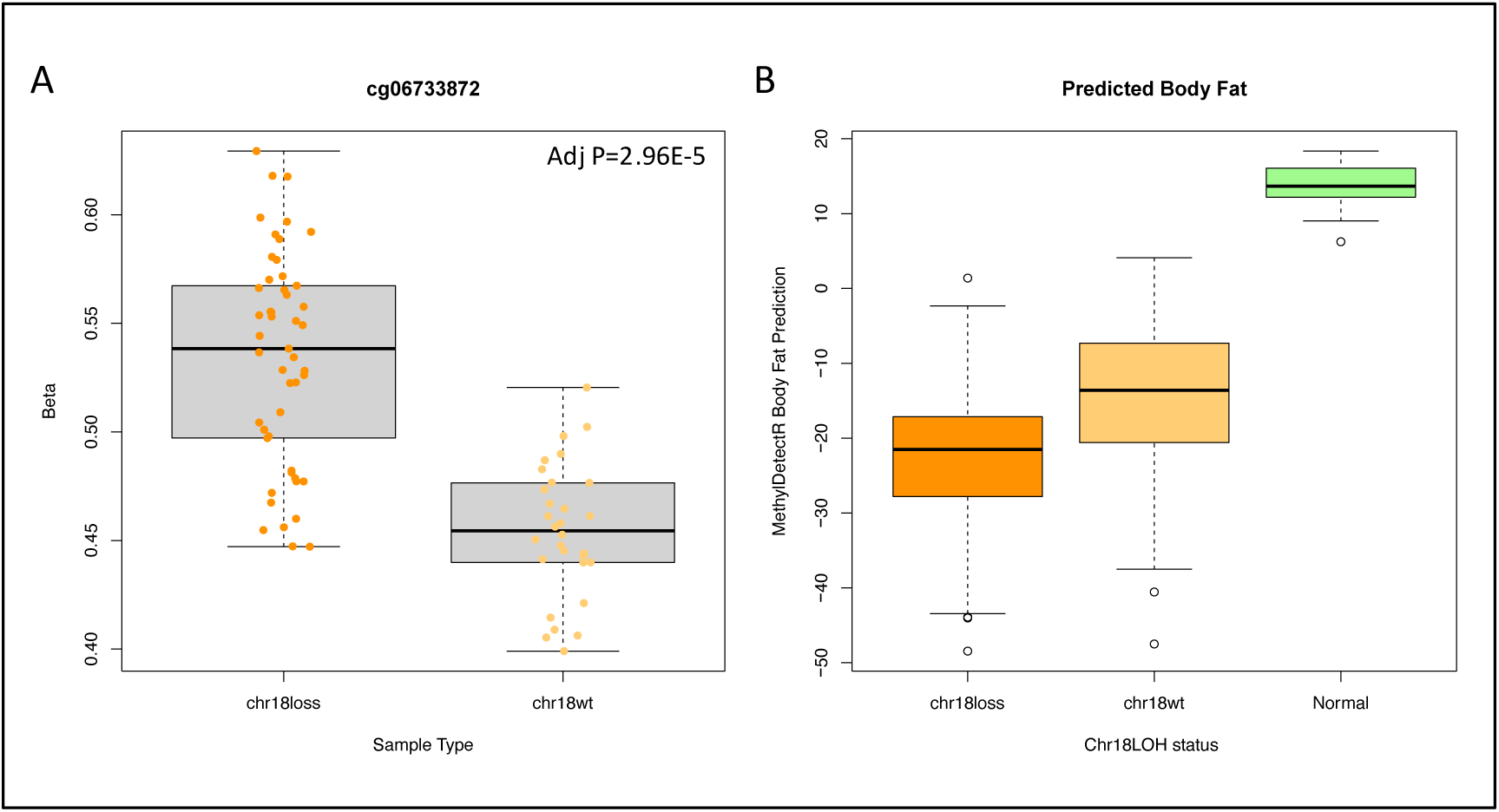
Differential methylation in comparison of tumours with any chromosome 18 LOH vs no LOH. (A) The top differentially methylated position between these comparison groups. Tumours containing chromosome 18 LOH had significantly higher methylation at this CpG site compared to tumours with wild type chromosome 18. (B) Predicted ‘body fat’ was significantly lower in chromosome 18 LOH tumours compared with chromosome 18 wild type tumours (p=0.00062), normal epithelia samples were not included in comparison and are just shown in plot for reference.

**Table 2:** Top 10 DMPs associated with chromosome 18 loss of heterozygosity in SI-NETs. Chr: chromosome, UCSC RefGene Name/Group: name of gene or gene element annotation as defined by the UCSC Genome Browser, according to the EPIC Illumina manifest.

| Probe | logFC | PVal | Bonf Adj PVal | UCSC RefGene Name | UCSC RefGene Group | Chr | Relation to CpG Island |
| --- | --- | --- | --- | --- | --- | --- | --- |
| cg06733872 | 0.07708657 | 4.47e-11 | 2.96e-05 | BDP1 | 3'UTR | 5 |  |
| cg09830678 | 0.03497408 | 1.94e-10 | 0.000128 | MPPE1 | 1stExon;5'UTR | 18 | Island |
| cg26280179 | -0.143416 | 4.05e-10 | 0.000268 |  |  | 10 |  |
| cg07788111 | -0.1846264 | 8.82e-10 | 0.000584 | LRRTM2;CTNNA1 | Body;TSS1500 | 5 |  |
| cg08048567 | -0.1657179 | 1.44e-09 | 0.000952 |  |  | 10 |  |
| cg01857824 | -0.1672307 | 1.66e-09 | 0.00110 | FYN | Body | 6 |  |
| cg11770586 | 0.01690181 | 2.69e-09 | 0.00178 | PSMG2; CEP76 | TSS1500;Body | 18 | Island |
| cg04167854 | -0.169423 | 5.45e-09 | 0.00361 | SERGEF | Body | 11 | N_Shelf |
| cg13158235 | -0.2033734 | 5.80e-09 | 0.00384 | DERA | Body | 12 |  |
| cg10543953 | -0.0777852 | 6.64e-09 | 0.00439 | LOC723972 | Body | 15 |  |

## Discussion

Recent investigations into somatic mutational landscape of multifocal SI-NETs have provided robust evidence that these tumours develop independently of each other, and that they display genetic heterogeneity within and between patients, but the few driver genes identified in these tumours to date suggests that epigenetic disruption is likely to be key in the evolution of these tumours (*8, 9, 11*). In this study, we have investigated genome-wide DNA methylation in multifocal SI-NETs for the first time. We assessed epigenetic ageing, differential methylation of genes and metabolic traits across multiple primary tumours, metastatic and matched normal tissue within a unique cohort of 11 patients diagnosed with metastatic multifocal SI-NETs.

The low abundance of the EC cells in matched normal epithelium samples has been a weakness of all investigations into the DNA methylation of SI-NETs to date, particularly in the identification of DMPs. In this study, we present a novel approach which combined these matched heterogeneous intestinal samples with methylation profile of a cultured EC-enriched cell line to triangulate the most biologically plausible tumour-associated methylation differences in multifocal SI-NETs. While this cell line includes multiple enteroendocrine cell types, it is substantially enriched for EC cells making it useful for refinement of the tumour-specific signal and reduction of the ‘noise’ in the signal caused by cellular differences between the tumours and the normal small intestinal epithelial tissue. While these DMPs were identified between tumour and normal small intestinal tissue, this approach will reduce the ‘falsepositive’ DMP signal driven by these cell composition differences. By excluding any methylation differences identified between EC cell-enriched cell line and normal tissue and focussing only on DMPs identified in comparison of primary tumours with both normal and cultured cell line, we have identified an SI-NET methylation signature which is specific to tumour cells and should be enriched for signals specific to the tumorigenic transition from normal EC cell to SI-NET tumour. This approach provides greater potential to identify relevant genes and pathways including potential therapeutic targets. Using this approach, 12,392 tumour-specific DMPs were identified which were enriched for neural pathways and genes relating to synaptic pathways. The most significant DMP mapped to the *SPTBN4* gene which contained 16 DMPs (Bonferroni adjusted P<0.05) in the refined analysis, indicating significant gene-wide methylation differences in the tumours. The biological importance of this gene in SI-NETs was also supported by the identification of significant differential expression of this gene between tumour and normal samples. This gene, ‘*spectrin beta, non-erythrocytic 4*’ is a protein coding gene which is overexpressed in the brain (*49*). It is associated with neurodevelopmental disorder with hypotonia, neuropathy and deafness and is related to the RET signalling pathway and cytokine signalling. The role of this gene in RET signalling is of particular relevance to SI-NETs as RET is a proto-oncogene which has previously been implicated in neuroendocrine carcinogenesis, as germline mutations in RET lead to the hereditary cancer syndrome, multiple endocrine neoplasia type 2 (MEN 2) (*50*).

As multifocal SI-NETs are slow-growing and generally remain small, to date it has been extremely challenging to assess the order in which tumours have developed within a patient. In this study, we have demonstrated that epigenetic clocks can be used to predict the order in which tumours developed, and we have determined that the Horvath Skin and Blood clock appears to perform this prediction most accurately. This was based on the assumption that normal intestinal tissue within these patients should cluster most closely with the chronological age of the patients. However, it is also important to consider that the adjacent normal tissue of a patient with multiple SI-NET tumours could possibly experience a degree of age acceleration caused by a field effect of the tumour. The cell of origin of SI-NETs is thought to be EC cells which occur at a low frequency within the intestinal tissue, which has been treated as the matched normal comparison for this cohort. As we know cell and tissue type have a huge impact on DNA methylation profiles (*48*), it is possible that the epigenetic age of purified EC cells may differ from the heterogeneous intestinal tissue investigated here. The variability seen in the predictions of tumour age across tumours from each patient provides support for the independent evolution theory of multifocal SI-NETs (*8, 9, 11*). Metastatic tumours appeared to retain the epigenetic age of the tumour from which they were derived. However, this was not statistically significant, possibly due to the small sample size and warrants further investigation in a larger cohort to better understand epigenetic ageing and how much it is influenced by the microenvironment of surrounding tissue versus tissue of origin. While the Skin and Blood clock appeared to perform best at estimating chronological age of normal tissue and discriminating development order in tumours out of 11 epigenetic clocks tested in this study, which we believe is due to its’ optimisation for epithelial tissue, it must be noted that this clock was not developed for use on cancer tissue.

For the first time, we have demonstrated systematic epigenetic alteration of key metabolic genes in the DNA methylation patterns of multifocal SI-NETs. Initially we identified this pattern through use of DNAm-score based prediction of several metabolic traits as implemented by the MethylDetectR tool. However, a weakness of this approach is that these scores were developed for use in blood, and to date have not been validated for application in cancer tissue. In the application to tumour samples performed in our study, we expect that this variation in DNAm score performance is reflective of the substantial differential methylation across genes associated with metabolism we observed in tumours. These changes are seen in the methylation levels of key metabolic genes when assessed independently of the MethylDetectR predictor. We hypothesise that these differences could be driven by nutrient restriction within the tumours, or due to increased metabolism of nutrients within tumours. The patterns of altered methylation identified in primary tumours compared with matched normal tissue were found to be more pronounced in metastatic tumours, and in tumours with chromosome 18 LOH. We believe this implies that these subgroups of tumours have some underlying difference in their metabolism or nutrient availability. As these patients are known from previous analyses to have primary tumours both with and without chromosome 18 LOH (*11*), this indicates that tumours within a single patient can have differences in their metabolic traits.

Chromosome 18 LOH is the most common genetic disruption in SI-NETs, and while patients with tumours harbouring this genomic alteration are observed to have better progression-free survival, the underlying mechanism is not understood. In a previous study of single SI-NETs, 27,464 chromosome 18 LOH associated DMPs were identified alongside 12 differentially expressed genes which were enriched for differential methylation (*21*). Waterfield *et al.* also identified three genes that were found to have differentially methylated regions that directly related to differential expression of the matched samples (*SCRT1*, *AMPD3* and *GRAMD2*) and in the current study each of these genes was found to contain differential methylation in comparison between tumours with and without chromosome 18 LOH (FDR p<0.05). This indicates shared epigenetic differences between tumours with chromosome 18 LOH whether they are single or multifocal tumours. Further work is needed to establish the epigenetic differences and similarities between single and multifocal SI-NETs.

## Conclusions

In the first study of epigenetic profiles in multifocal SI-NETs to date, we have identified tumour-specific differential methylation, epigenetic age acceleration and tumour-associated metabolic differences reflected in the epigenome for the first time. Methylation differences associated with the development of SI-NETs were identified using a novel approach, combining data from matched patient-derived small intestinal tissue and cultured cells enriched for the EC cells, to reduce the impact of cell heterogeneity on findings. From this analysis we have identified several neural-specific genes which are consistently differentially methylated across multifocal SI-NETs and which may provide candidates for therapeutic intervention following further research. We have also demonstrated the use of epigenetic clocks for assessing the order of occurrence of multifocal SI-NETs within a patient, which provides an opportunity to investigate how these tumours develop and to assess if there are any distinct molecular features of the initial tumour, which might lead to development of other tumours in the surrounding tissue. We have also identified epigenetic alterations in key metabolic genes, which we believe is reflecting metabolic differences in SI-NETs which are more pronounced in metastatic tumours and tumours with chromosome 18 LOH.

## Acknowledgements

The authors would like to acknowledge the UCL Genomics facility for generating the DNA methylation data. The authors would also like to thank the patients and their families for their contribution to this study.

## Author Contributions

Conceptualization, E.N., C.T., and A.P.W.;

Formal Analysis, A.P.W. and N.Ma.;

Investigation, A.P.W, N.Ma., S.E., N.Me. and C.C;

Writing—Original Draft, A.P.W.;

Writing—Review & Editing, C.T., N.Me., C.C., S.E., M.S., P.Y. and N.Ma.;

Funding Acquisition, E.N., C.T.;

Resources, E.N., H.V. and P.D.;

Supervision, C.T. and N.M.

All authors read and approved the final manuscript.

## Supplementary Figures and Tables

**Supplementary Table 1:** Summary of samples included in the study for each patient. All patients had multiple primary tumours (ranging 2-16), a subset had metastatic tumours (ranging 1-2) and a subset had matched normal epithelium samples available.

| Patient ID | Normal Epithelia | Primary | Metastases | Total EPIC datasets |
| --- | --- | --- | --- | --- |
| P744 | 1 | 3 | 2 | **6** |
| P760 | 1 | 3 | 0 | **4** |
| P772 | 1 | 2 | 2 | **5** |
| P787 | 1 | 3 | 0 | **4** |
| P825 | 0 | 3 | 1 | **4** |
| P848 | 1 | 11 | 2 | **14** |
| P850 | 1 | 9 | 1 | **11** |
| P852 | 1 | 16 | 1 | **18** |
| P876 | 1 | 8 | 1 | **10** |
| P947 | 1 | 10 | 0 | **11** |
| P952 | 0 | 11 | 2 | **13** |

**Supplementary Figure 1:**
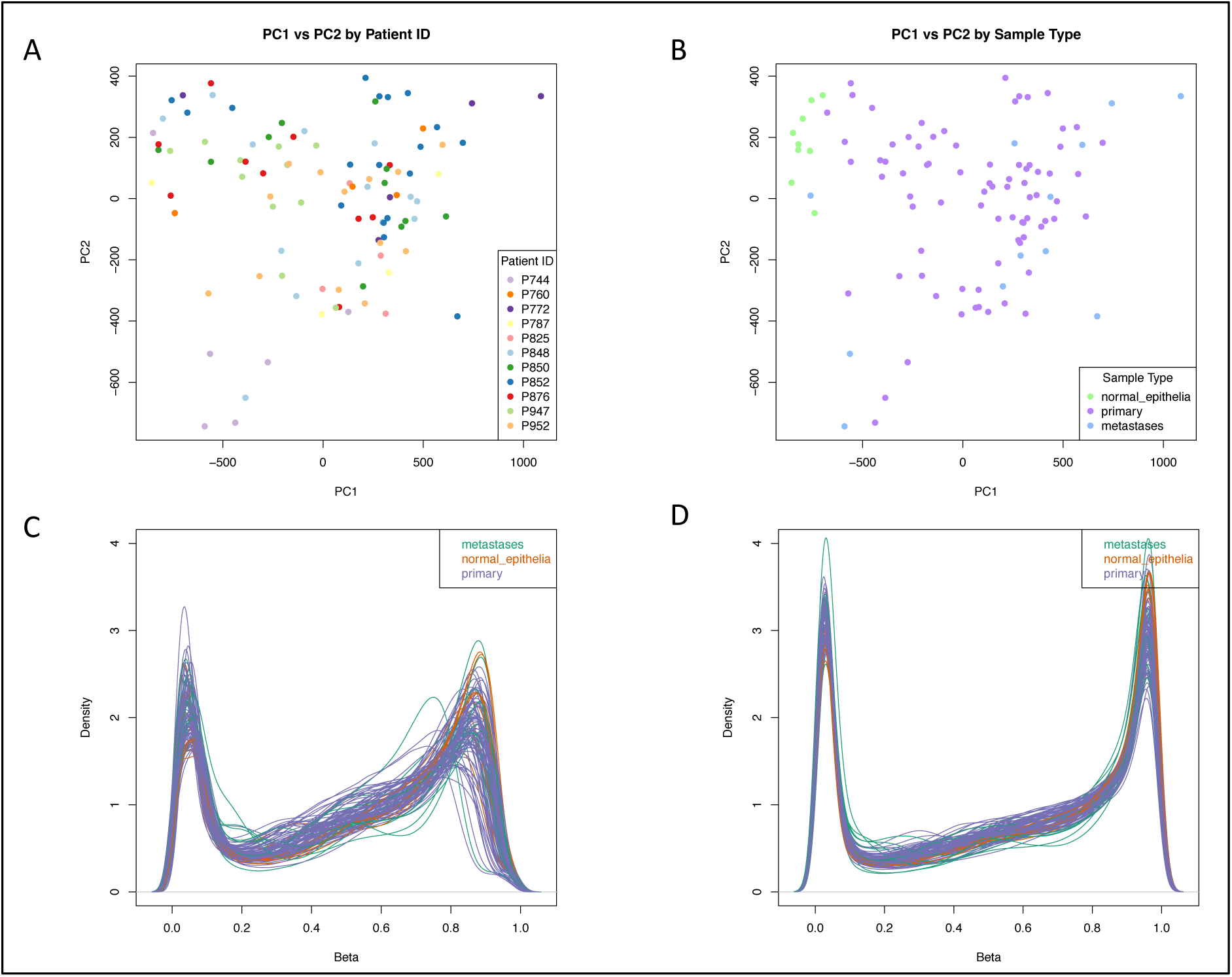
Quality assessment of dataset. Principal components analysis demonstrated that samples did not cluster by individual (A), but did cluster by sample type (B) when comparing principal components 1 and 2. The beta distributions for all samples (n=100) was good in the raw dataset (C) and was improved in the normalised dataset (D).

**Supplementary Figure 2:**
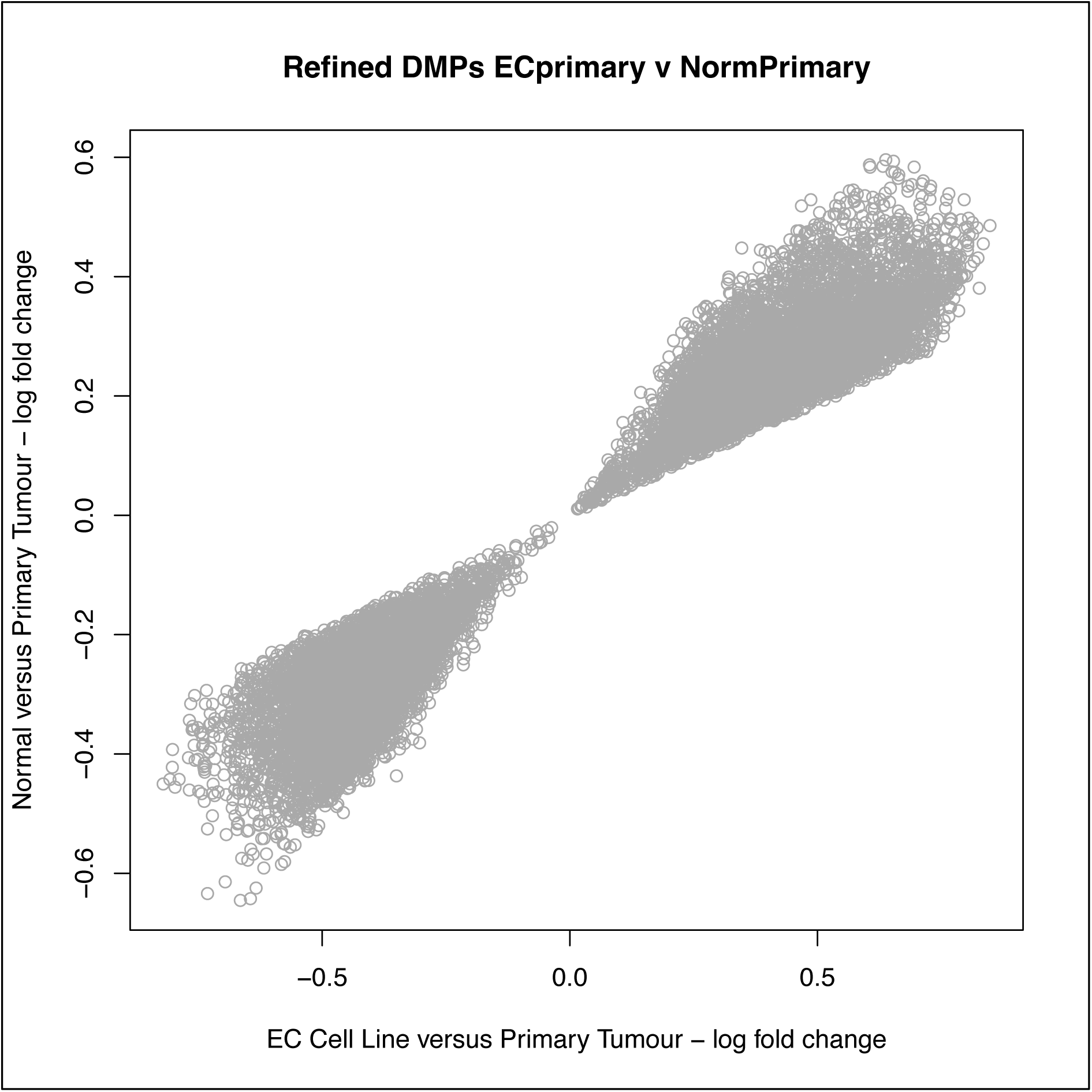
Direction of overlapping DMPs in tumour-related methylation analysis. Scatter plot demonstrating that all DMPs which overlapped between ‘EC cells versus Primary Tumour DMPs’ and ‘Normal versus Primary Tumour DMPs’, but were not significant in the ‘Normal vs EC cell DMPs’ were all differentially methylated in the same direction. Ie, sites were consistently hypermethylated or hypomethylated in tumours, whether they were compared with normal epithelia or EC cell line.

***Supplementary Table 2: List of refined tumour-associated DMPs (attached separately)****. Full list of 12,392 tumour-associated DMPs identified during analysis of SI-NET tumours, following refinement using an EC cell-enriched cell line*.

**Supplementary Table 3:** Top 20 pathways identified from gene ontology analysis of significant tumour-associated DMPs. CC: cellular component, BP: biological process, N: number of genes in the GO term, DE: number of genes that are differentially methylated, P.DE: p-value for over-representation of the GO term, FDR: false discovery rate adjusted p-value of pathway

| GO Pathway | Ontology | Term | N | DE | P.DE | FDR |
| --- | --- | --- | --- | --- | --- | --- |
| GO:0045202 | CC | synapse | 1289 | 507 | 2.10E-09 | 3.93E-05 |
| GO:0030054 | CC | cell junction | 1999 | 743 | 3.47E-09 | 3.93E-05 |
| **GO:0010646** | BP | regulation of cell communication | 3470 | 1122 | 4.38E-08 | 0.00031643 |
| **GO:0023051** | BP | regulation of signaling | 3508 | 1133 | 5.72E-08 | 0.00031643 |
| **GO:0043005** | CC | neuron projection | 1297 | 491 | 7.88E-08 | 0.00031643 |
| **GO:0042995** | CC | cell projection | 2227 | 780 | 8.90E-08 | 0.00031643 |
| **GO:0007267** | BP | cell-cell signaling | 1687 | 591 | 1.08E-07 | 0.00031643 |
| **GO:0120025** | CC | plasma membrane bounded cell projection | 2128 | 747 | 1.12E-07 | 0.00031643 |
| **GO:0005886** | CC | plasma membrane | 5093 | 1503 | 2.13E-06 | 0.00485538 |
| **GO:0030029** | BP | actin filament-based process | 776 | 309 | 2.21E-06 | 0.00485538 |
| **GO:0071944** | CC | cell periphery | 5206 | 1537 | 2.36E-06 | 0.00485538 |
| **GO:0051049** | BP | regulation of transport | 1750 | 580 | 5.06E-06 | 0.00953638 |
| **GO:0007268** | BP | chemical synaptic transmission | 696 | 274 | 5.97E-06 | 0.00964327 |
| **GO:0098916** | BP | anterograde trans-synaptic signaling | 696 | 274 | 5.97E-06 | 0.00964327 |
| **GO:0098794** | CC | postsynapse | 607 | 254 | 6.77E-06 | 0.01021612 |
| **GO:0008092** | MF | cytoskeletal protein binding | 938 | 359 | 8.41E-06 | 0.01189747 |
| **GO:0032879** | BP | regulation of localization | 2751 | 879 | 1.05E-05 | 0.01403586 |
| **GO:0051046** | BP | regulation of secretion | 674 | 240 | 1.16E-05 | 0.01458469 |
| **GO:0099537** | BP | trans-synaptic signaling | 703 | 275 | 1.37E-05 | 0.01636757 |
| **GO:0030036** | BP | actin cytoskeleton organization | 680 | 272 | 1.72E-05 | 0.01943346 |

**Supplementary Figure 3:**
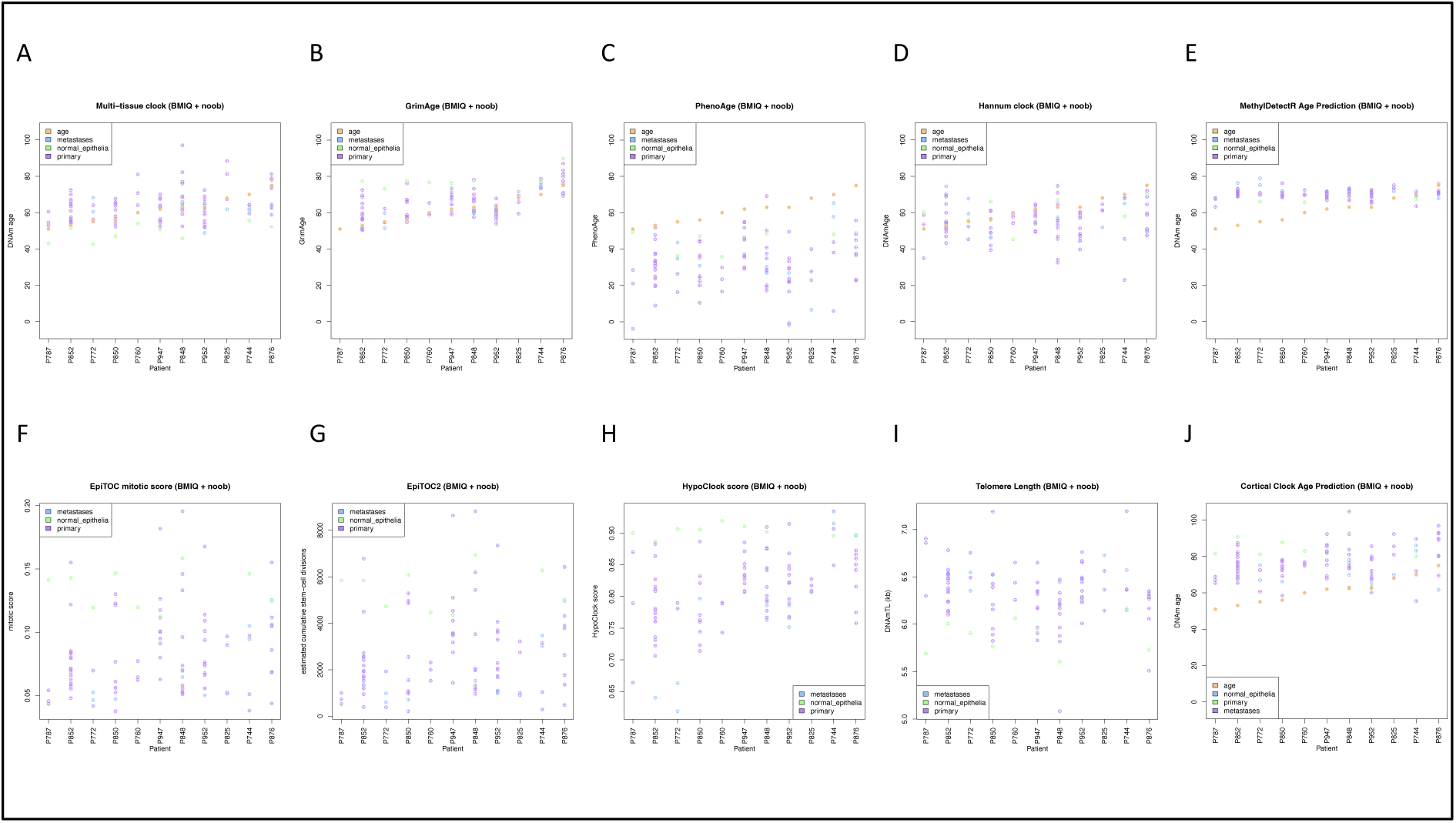
Comparison of predictions from ten epigenetic clocks in multifocal SI-NETs. Epigenetic clock predictions are shown for normal (green), primary tumours (purple) and metastatic tumours (blue), with chronological age indicated by the orange points in plots where this is appropriate (ie, in clocks whose output is in years). Patients are indicated in order of chronological age.

**Supplementary Figure 4:**
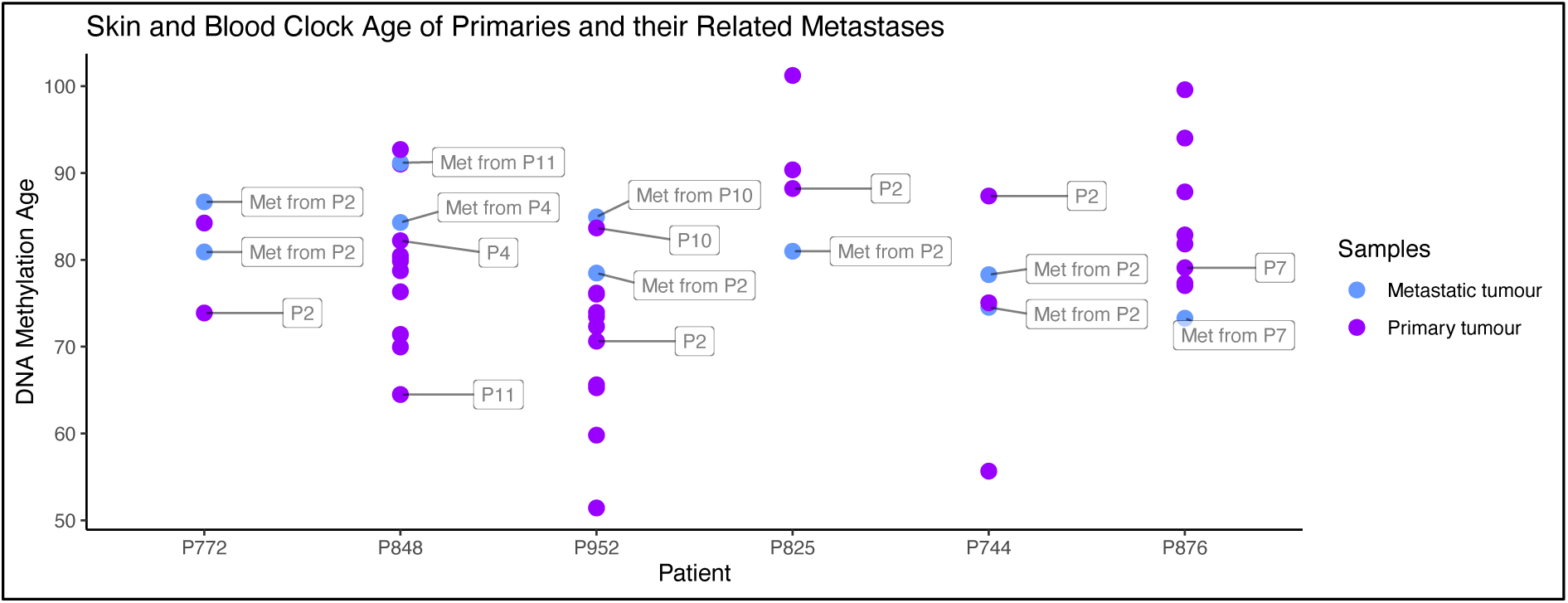
Skin and Blood clock predictions of metastatic tumours and the metastases they derive from.

## References

1. J. Hallet et al., Exploring the rising incidence of neuroendocrine tumors: a population-based analysis of epidemiology, metastatic presentation, and outcomes. Cancer 121, 589–597 (2015).

2. K. Y. Bilimoria et al., Small bowel cancer in the United States: changes in epidemiology, treatment, and survival over the last 20 years. Ann Surg 249, 63–71 (2009).

3. A. Dasari et al., Trends in the Incidence, Prevalence, and Survival Outcomes in Patients With Neuroendocrine Tumors in the United States. JAMA Oncol 3, 1335–1342 (2017).

4. J. Darbà, A. Marsà, Exploring the current status of neuroendocrine tumours: a population-based analysis of epidemiology, management and use of resources. BMC Cancer 19, 1226 (2019).

5. A. Gangi et al., Multifocality in Small Bowel Neuroendocrine Tumors. J Gastrointest Surg 22, 303–309 (2018).

6. A. B. Choi et al., Is Multifocality an Indicator of Aggressive Behavior in Small Bowel Neuroendocrine Tumors? Pancreas 46, 1115–1120 (2017).

7. R. K. Yantiss, R. D. Odze, F. A. Farraye, A. E. Rosenberg, Solitary versus multiple carcinoid tumors of the ileum: a clinical and pathologic review of 68 cases. Am J Surg Pathol 27, 811–817 (2003).

8. Z. Zhang et al., Patterns of chromosome 18 loss of heterozygosity in multifocal ileal neuroendocrine tumors. Genes Chromosomes Cancer 59, 535–539 (2020).

9. E. Elias et al., Independent somatic evolution underlies clustered neuroendocrine tumors in the human small intestine. Nat Commun 12, 6367 (2021).

10. T. M. Katona et al., Molecular evidence for independent origin of multifocal neuroendocrine tumors of the enteropancreatic axis. Cancer Res 66, 4936–4942 (2006).

11. N. Mäkinen et al., Whole genome sequencing reveals the independent clonal origin of multifocal ileal neuroendocrine tumors. Genome Med 14, 82 (2022).

12. M. S. Banck et al., The genomic landscape of small intestine neuroendocrine tumors. J Clin Invest 123, 2502–2508 (2013).

13. P. Priestley et al., Pan-cancer whole-genome analyses of metastatic solid tumours. Nature 575, 210–216 (2019).

14. M. H. Kulke et al., High-resolution analysis of genetic alterations in small bowel carcinoid tumors reveals areas of recurrent amplification and loss. Genes Chromosomes Cancer 47, 591–603 (2008).

15. S. Kytölä et al., Comparative genomic hybridization identifies loss of 18q22-qter as an early and specific event in tumorigenesis of midgut carcinoids. Am J Pathol 158, 1803–1808 (2001).

16. R. M. Löllgen, O. Hessman, E. Szabo, G. Westin, G. Akerström, Chromosome 18 deletions are common events in classical midgut carcinoid tumors. Int J Cancer 92, 812–815 (2001).

17. J. L. Cunningham et al., Common pathogenetic mechanism involving human chromosome 18 in familial and sporadic ileal carcinoid tumors. Genes Chromosomes Cancer 50, 82–94 (2011).

18. J. M. Francis et al., Somatic mutation of CDKN1B in small intestine neuroendocrine tumors. Nat Genet 45, 1483–1486 (2013).

19. J. Crona et al., Somatic Mutations and Genetic Heterogeneity at the CDKN1B Locus in Small Intestinal Neuroendocrine Tumors. Ann Surg Oncol 22 Suppl 3, S1428–1435 (2015).

20. A. Karpathakis et al., Prognostic Impact of Novel Molecular Subtypes of Small Intestinal Neuroendocrine Tumor. Clin Cancer Res 22, 250–258 (2016).

21. S. Waterfield et al., Chromosome 18 loss of heterozygosity in small intestinal neuroendocrine tumours: Multi-omic and tumour composition analyses. Neuroendocrinology, (2023).

22. A. Dobin et al., STAR: ultrafast universal RNA-seq aligner. Bioinformatics 29, 15–21 (2013).

23. B. Li, C. N. Dewey, RSEM: accurate transcript quantification from RNA-Seq data with or without a reference genome. BMC Bioinformatics 12, 323 (2011).

24. M. I. Love, W. Huber, S. Anders, Moderated estimation of fold change and dispersion for RNA-seq data with DESeq2. Genome Biol 15, 550 (2014).

25. M. J. Aryee et al., Minfi: a flexible and comprehensive Bioconductor package for the analysis of Infinium DNA methylation microarrays. Bioinformatics 30, 1363–1369 (2014).

26. Y. Tian et al., ChAMP: updated methylation analysis pipeline for Illumina BeadChips. Bioinformatics 33, 3982–3984 (2017).

27. Z. Xu, L. Niu, L. Li, J. A. Taylor, ENmix: a novel background correction method for Illumina HumanMethylation450 BeadChip. Nucleic Acids Res 44, e20 (2016).

28. M. E. Ritchie et al., limma powers differential expression analyses for RNA-sequencing and microarray studies. Nucleic Acids Res 43, e47 (2015).

29. B. Phipson, J. Maksimovic, A. Oshlack, missMethyl: an R package for analyzing data from Illumina’s HumanMethylation450 platform. Bioinformatics 32, 286–288 (2016).

30. A. E. Teschendorff et al., A beta-mixture quantile normalization method for correcting probe design bias in Illumina Infinium 450 k DNA methylation data. Bioinformatics 29, 189–196 (2013).

31. J. P. Fortin, T. J. Triche, K. D. Hansen, Preprocessing, normalization and integration of the Illumina HumanMethylationEPIC array with minfi. Bioinformatics 33, 558–560 (2017).

32. W. Zhou, P. W. Laird, H. Shen, Comprehensive characterization, annotation and innovative use of Infinium DNA methylation BeadChip probes. Nucleic Acids Res 45, e22 (2017).

33. P. N. P. Singh et al., Transcription factor dynamics, oscillation, and functions in human enteroendocrine cell differentiation. Cell Stem Cell, (2024).

34. J. Maksimovic, A. Oshlack, B. Phipson, Gene set enrichment analysis for genome-wide DNA methylation data. Genome Biol 22, 173 (2021).

35. P. Geeleher et al., Gene-set analysis is severely biased when applied to genome-wide methylation data. Bioinformatics 29, 1851–1857 (2013).

36. S. Horvath, DNA methylation age of human tissues and cell types. Genome Biol 14, R115 (2013).

37. S. Horvath et al., Epigenetic clock for skin and blood cells applied to Hutchinson Gilford Progeria Syndrome and. Aging (Albany NY) 10, 1758–1775 (2018).

38. A. T. Lu et al., DNA methylation GrimAge strongly predicts lifespan and healthspan. Aging (Albany NY) 11, 303–327 (2019).

39. M. E. Levine et al., An epigenetic biomarker of aging for lifespan and healthspan. Aging (Albany NY) 10, 573–591 (2018).

40. G. Hannum et al., Genome-wide methylation profiles reveal quantitative views of human aging rates. Mol Cell 49, 359–367 (2013).

41. R. F. Hillary, R. E. Marioni, MethylDetectR: a software for methylation-based health profiling. Wellcome Open Res 5, 283 (2020).

42. Z. Yang et al., Correlation of an epigenetic mitotic clock with cancer risk. Genome Biol 17, 205 (2016).

43. A. E. Teschendorff, A comparison of epigenetic mitotic-like clocks for cancer risk prediction. Genome Med 12, 56 (2020).

44. A. T. Lu et al., DNA methylation-based estimator of telomere length. Aging (Albany NY) 11, 5895–5923 (2019).

45. G. L. Shireby et al., Recalibrating the epigenetic clock: implications for assessing biological age in the human cortex. Brain 143, 3763–3775 (2020).

46. E. C. D. Foundation. (https://dnamage.genetics.ucla.edu/home), vol. 2021.

47. R. F. Hillary, R. E. Marioni. (https://shiny.igmm.ed.ac.uk/Calculate_Your_Scores/), vol. 2021.

48. N. Loyfer et al., A DNA methylation atlas of normal human cell types. Nature 613, 355–364 (2023).

49. L. Fagerberg et al., Analysis of the human tissue-specific expression by genome-wide integration of transcriptomics and antibody-based proteomics. Mol Cell Proteomics 13, 397–406 (2014).

50. Y. Murakumo, M. Jijiwa, N. Asai, M. Ichihara, M. Takahashi, RET and neuroendocrine tumors. Pituitary 9, 179–192 (2006).

